# Neural mechanisms of object prioritization in vision

**DOI:** 10.1101/2025.01.09.632156

**Authors:** Damiano Grignolio, Sreenivasan Meyyappan, Joy Geng, George R. Mangun, Clayton Hickey

## Abstract

Selective attention is widely thought to be sensitive to visual objects. This is commonly demonstrated in cueing studies, which show that when attention is deployed to a known target location that happens to fall on a visual object, responses to targets that unexpectedly appear at other locations on that object are faster and more accurate, as if the object in its entirety has been visually prioritized. However, this notion has recently been challenged by results suggesting that putative object-based effects may reflect the influence of hemifield anisotropies in attentional deployment, or of unacknowledged influences of perceptual complexity and visual clutter. Studies employing measures of behaviour provide limited opportunity to address these challenges. Here, we used EEG to directly measure the influence of task-irrelevant objects on the deployment of visual attention. We had participants complete a simple visual cueing task involving identification of a target that appeared at either a cued location or elsewhere. Throughout each experimental trial, displays contained task-irrelevant rectangle stimuli that could be oriented horizontally or vertically. We derived two cue-elicited indices of attentional deployment–lateralized alpha oscillations and the ADAN component of the event-related potential–and found that these were sensitive to the otherwise irrelevant orientation of the rectangles. Our results demonstrate that the allocation of visual attention is influenced by objects boundaries, supporting models of object-based attentional prioritization.

## Introduction

Adaptive behaviour relies on identifying useful objects in our environment. Accordingly, sensory and cognitive systems, including visual attention, express object-based organization (<u>Scholl, 2001</u>; <u>Duncan, 1984</u>). This appears both in studies of patients and healthy controls. In patients, parietal lesions degrade the ability to attend to multiple objects simultaneously (<u>Luria, 1959</u>; <u>Coslett, & Saffran, 1991</u>) and bias attention to the ipsi-lesional side of objects (<u>Walker, 1995</u>). In healthy controls, attention is more efficiently deployed to stimuli that fall on a cued object rather than outside that object (<u>Chen et al., 2012</u>).

This latter instantiation of object prioritization is commonly studied using the two-rectangle task (Egly, Driver, & Rafal, 1994; Moore, Yantis, & Vaughan, 1998). In this paradigm, attention is cued - either by an exogenous cue such as luminance onset (eg. Egly, Driver, & Rafal, 1994) or by an endogenous cue such as a central arrow (eg. Abrams & Law, 2000, Exp. 2; Chen & Cave, 2008) - to an endpoint of one of two rectangles rendered on the computer screen (Figure 1). Responses are faster and more accurate when attention is invalidly cued to an endpoint that happens to fall within the same rectangle as the cued location, as compared to when the target appears at any other uncued position.

**Figure 1.**
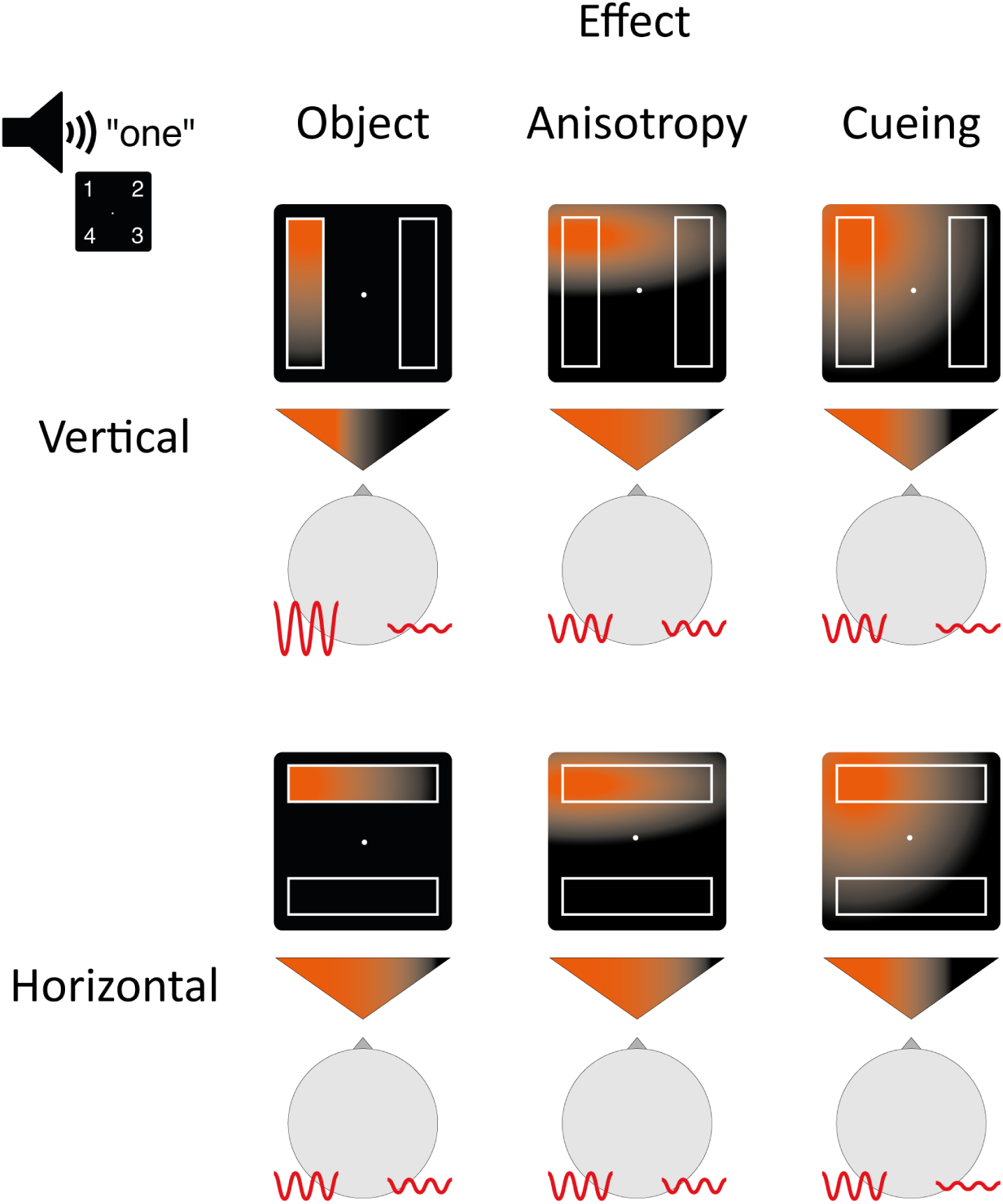
Attention deployment in horizontal and vertical conditions under the influence of three effects: object-based attention (object), attentional anisotropy (anisotropy) and spatial selection (cueing). An auditory cue indicates position one (upper-left corner). The orange gradient represents the distribution of attention across the screen following the cue, while the gradient within the triangles illustrates the direction or lateralization of attention. The red lines depict alpha oscillations ipsilateral and contralateral to the cued position, as well as their power. As illustrated, attentional anisotropy and cueing do not predict an effect on alpha lateralization across horizontal and vertical rectangle conditions, but object-based attention does predict such an effect. Similar expectations can be derived for the ADAN. Importantly, these hypothesized effects are not exclusive and may combine simultaneously.

This has been interpreted as reflecting the automatic spreading of spatial attention along the contours of the cued object, reflecting a role for attention in object completion and other Gestalt principles of visual perception (Davis & Driver, 1997; Cohen et al., 2015; Watson & Kramer, 1999). However, recent studies have raised challenges to this idea (Reppa, Schmidt, & Leek, 2012; Francis & Thunnell, 2022).

First, object prioritization appears to be contingent on cognitive strategy. For example, Shomstein and Yantis (2002) found that non-targets on the same object as targets had no impact on performance unless there was uncertainty about target location. This is difficult to reconcile with the notion that object prioritization is closely involved in low-level perception, which should not be sensitive to task set (Shomstein, 2012). Second, object effects in the two-rectangle task are worryingly sensitive to rectangle orientation, emerging with more strength when the rectangles are horizontal (Al-Janabi & Greenberg, 2016; Chen & Cave, 2019; Francis & Thunnell, 2022; Pilz, Roggeveen, Creighton, Bennett, & Sekuler, 2012). Performance of many visual tasks is known to be better when stimuli appear across the horizontal meridian (Corballis & Roldan, 1975; Carrasco, Giordano, & McElree, 2004; Corbett & Carrasco, 2011), raising the possibility that putative object prioritization might instead reflect an influence of hemifield anisotropy (Pilz, Roggeveen, Creighton, Bennett, & Sekuler, 2012; Barnas & Greenberg, 2024). Third, interpretation of object-based effects is complicated by reporting bias. A meta-analysis of 37 behavioural studies found that 19 were unlikely to replicate with the original sample size (Francis & Thunnell, 2022) and a large-sample study failed to replicate the effect (Pilz et al., 2012, Exp. 2).

Motivated by these and other findings, Rosenholtz (2024) has recently challenged the notion that behavioural measures of object prioritization reflect attention at all, noting that when cue and target appear on different objects, the edges of both objects intervene. This could generate a small cost to target resolution through visual crowding, and this could be exacerbated if participants move their eyes to bring the visual clutter closer to foveal vision - which is likely, given that studies of object prioritization commonly employ cue validity of >70% without monitoring fixation. In line with this, the effect of object prioritization reverses when visual complexity is introduced into the space between same-object locations (Chen, Cave, Basu, Suresh, & Wiltshire, 2020).

It is therefore unclear if attention is necessarily sensitive to the presence of visual objects at all. This uncertainty is caused in part by a reliance in the literature on inference from behaviour. On one hand, issues like stimulus crowding, complexity, and hemifield anisotropies can create costs in the behavioural response to targets that are difficult to distinguish from effects of object prioritization. On the other, object prioritization may simply degrade over time (Lou, Lorist, & Pilz, 2023), such that small, absent, or even reversed effects at the time of target onset do not mean that earlier deployment of attention was unaffected. Here, we resolve this ambiguity by deriving indices of attention from human EEG, allowing us to directly measure the impact of task-irrelevant objects on the deployment of attention.

## Methods

We had participants complete a variant of the two-rectangle paradigm where target location was endogenously cued by a spoken word identifying a screen location (Goldsmith & Yeari, 2003). We recorded EEG while participants completed this task and derived two indices of attentional deployment from this signal: lateralized posterior EEG alpha power (Worden et al., 2000), and an ERP component known as the anterior directing-attention negativity (ADAN), which reflects activation of frontal brain structures involved in the strategic control of attention (Eimer et al, 2002).

Occipital alpha oscillations are known to be modulated by the deployment of attention in retinotopic space (<u>Popov et al., 2019</u>). In particular, when attention is directed to a lateral position, alpha oscillations originating from contralateral posterior cortex exhibit lower amplitude compared to those originating ipsilaterally. This is thought to reflect the preparatory downregulation of ongoing inhibitory activity, such that visual cortex contralateral to the cued location becomes broadly more responsive to stimulus inputs (Klimesch et al., 2007; Jensen and Mazaheri, 2010; Foxe and Snyder, 2011).

In contrast, the ADAN emerges over lateral frontal brain areas. It is evoked by endogenous attention-directing cues and is associated with activation of cortical areas involved in the control and voluntary deployment of spatial attention (<u>Praamstra 2005</u>, <u>Eimer et al. 2002</u>; <u>Hopf and Mangun, 2000</u>). It typically emerges 300-500 ms after cue onset and is characterized by a negative deflection in the ERP waveform that is more pronounced over frontal and central scalp regions contralateral to the focus of attention (see <u>Zani et al., 2023</u>; <u>Holmes et al., 2010</u>; <u>Seiss et al., 2007</u>; <u>Eimer et al., 2002</u>; <u>Hopf & Mangun, 2000</u>; <u>Nobre et al., 2000</u>; <u>van Velzen, Forster, & Eimer, 2002; Yamaguchi et al., 1995b</u>).

As depicted in Figure 1, we hypothesized that lateralized alpha activity and the ADAN would differ based on the orientation of the irrelevant rectangles used in our task. When the rectangles are oriented vertically, each lies entirely within a single visual hemifield. If object-based attention causes selection of the cued object in this circumstance, attention will be strongly lateralized because the effects of spatial cuing and object-based attention will align. This should accordingly create strong lateralization of occipital EEG alpha activity and the ADAN. Conversely, when the rectangles are oriented horizontally, each rectangle appears in both visual hemifields. If object-based attention causes selection of the cued object in this circumstance, attention will be less lateralized because the effects of spatial cuing and object-based attention will be incongruent. This should create weaker lateralization of occipital alpha and the ADAN.

Unlike behavioral studies of object-based attention, which compare reaction times for validly versus invalidly cued targets on the same versus different objects, our critical comparison focuses solely on the vertical versus horizontal rectangle conditions. The dependent measures are the post-cue/pre-target EEG alpha activity and ADAN, which reflect the strength of anticipatory lateralization of spatial attention before the target appears.

### Participants

Thirty participants (6 males, 24 females; mean age 23.13 years ± 4,12 years SD; 6 left-handed) were recruited from the University of California, Davis, community. All had normal or corrected-to-normal vision and received compensation for their participation (mean compensation $45). All participants gave informed written consent, and the study procedure was approved by the local institutional review board of the University of California, Davis.

### Stimuli, Experimental Task, and Apparatus

Figure 2 depicts the stimuli and timing of an experimental trial. Each trial began with an auditory cue that indicated one of 4 screen locations where a subsequent target might appear, each located 4.4° visual angle from the screen center. The cue constituted a male voice with a standard British accent pronouncing one of four numbers: ‘one’, ‘two’, ‘three’, or ‘four’. Each number took 300 ms to pronounce and corresponded to a specific corner on the screen of an LCD computer monitor (57 cm x 39 cm; 120 Hz refresh rate), with ‘one’ designating the upper left corner and proceeding clockwise. The mapping between numbers and screen locations remained consistent across all participants and the cues were generated using Audacity (v3.2.4, Audacity Team, 2021).

**Figure 2.**
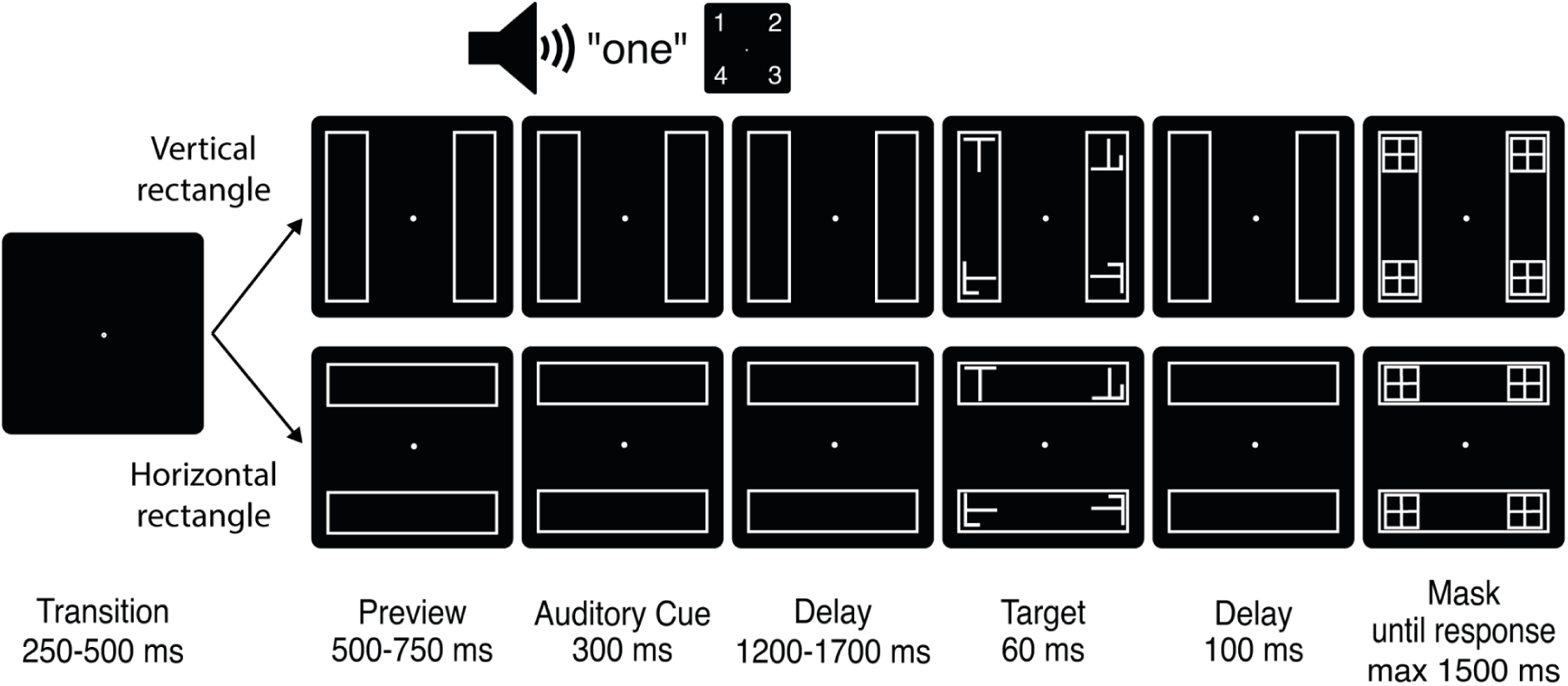
Task description: A fixation dot was presented for a variable duration of 250 to 500 ms, followed by a preview screen displaying the two rectangles for a variable duration of 500 to 750 ms. A 300-ms auditory cue then identified one of the four rectangle endpoints where the target was 70% likely to appear. The target and nontargets subsequently appeared after a delay of 1500 to 2000 ms. The target remained on the screen for 60 ms, followed by a 100 ms interval before the four potential target locations were masked. The mask sustained for 1500 ms or until response. Left index finger response indicated that the target was an ‘L’ and right index finger response indicated that the target was a ‘T’.

The target was either the character ‘L’ or ‘T’, and participants were required to report the identity of this stimulus by pressing the corresponding ‘L’ or ‘T’ key on a standard QWERTY computer keyboard with the left or right index finger. Nontargets appeared at the 3 screen locations where the target did not appear, and these were generated by superimposing the two target letters and applying a random rotation of 90°, 180°, or 270°. The target and nontarget stimuli subtended a visual angle of 1.2° x 0.9°, and appeared for 60 ms before being masked 100 ms later (Figure 2). The mask remained present until either the participant made a response or 1500 ms had passed, at which time the trial concluded.

In two-thirds of trials, the target appeared at the cued location. In the remaining trials, the target appeared with equal probability at the uncued end of the <u>cued</u> rectangle or at the nearest end of the <u>uncued</u> rectangle. For example, if position “one” (upper left) was cued, the target had two-thirds probability of appearing in position “one”, and otherwise appeared at either position “two” or “four” with equal probability.

Throughout each experimental trial, two rectangle stimuli were present on the screen (10° x 2.5° visual angle). The rectangles were created such that each of the four potential target locations lay at a rectangle endpoint. The orientation of these rectangles varied randomly from trial to trial, and was either vertical (such that left and right hemifield stimulus locations appeared on the same rectangle objects) or horizontal (such that top and bottom stimulus locations appeared on the same rectangle objects).

The experiment was programmed using Python and JavaScript in conjunction with the Opensesame software package (v3.3.6, Mathôt et al., 2012). The first participant in the experiment completed 720 trials, while the remaining participants completed 960 trials each. Detailed instructions emphasizing the importance of both speed and accuracy were provided to participants at the beginning of the experiment. Participants underwent a practice phase to familiarize themselves with the cue-location association. This phase involved a point-and-click task in which participants identified the location indicated by the auditory cue. Practice continued until participants achieved an average reaction time below 1300 ms and at least 80% accuracy. At the beginning of each practice trial, the mouse pointer was positioned at the center of the screen.

### EEG Recording and Pre-processing

EEG data were continuously sampled at a rate of 1 kHz using a Neuroscan SynAmps 2 amplifier and sintered Ag/AgCl ActiCap Snap active electrodes (Brain Products GmBH). Electrodes were placed at 64 scalp locations according to a 10-10 montage (Oostenveld & Praamstra, 2001). Two additional electrodes were placed one centimeter lateral to the external canthi of each eye to measure the horizontal electrooculogram (HEOG), and two further electrodes were placed 1 cm above and below the center of the left eye to measure the vertical electrooculogram (VEOG) and blink potentials. Signals were referenced to FCz during recording.

The data were digitally downsampled to 500 Hz. A high-pass filter with a cutoff frequency of 0.01 Hz was applied to remove low-frequency drift (Acunzo, MacKenzie, & van Rossum, 2012). The "pop_cleanline" function from the EEGLAB toolbox (Delorme & Makeig, 2004) for MATLAB was used to eliminate interference from line noise (60 Hz). The EEG signal was subsequently re-referenced to the average signal of all scalp electrodes, leaving the EOG channels unaffected. To identify and remove artifacts, independent component analysis (ICA; Bell & Sejnowski, 1995) was conducted on a copy of the EEG dataset that had been high-pass filtered at 1 Hz, with ICA weights then copied and associated with the original dataset.

Artifact rejection was performed using a combination of automatic and manual procedures. Muscle activity, electrical noise, eye blinks, and other noise-associated components were first automatically labeled using the *ICLabel* classifier (<u>Pion-Tonachini, 2019</u>), with labels subsequently confirmed via visual inspection. Trials with eye movements were marked for rejection based on two measures. First, we applied an absolute signal deviation threshold of 20 **μ**V to the horizontal electrooculogram (HEOG) in the interval 0 to 800 ms after the cue. Second, we visually identified the ICA component reflecting horizontal eye movements, applying a subject-tailored absolute threshold to this signal to identify contaminated trials. The results from both approaches largely overlapped and trials identified as containing eye movements via either criteria were first individually inspected and then rejected from further analysis (2.28% of trials +/- 1.8% SD were discarded from either time window). Variance associated with artifactual and noise-associated ICA components, including residual variance stemming from eye movements, was subsequently removed from the data.

### Time-frequency Analysis

Time-frequency representations were calculated using the *newtimef* function in the EEGLAB toolbox (Delorme & Makeig, 2004). This involved convolution of the EEG signal with a series of complex Morlet wavelets estimating 40 linear-spaced frequencies from 5.9 Hz to 40.0 Hz, with wavelets sampling from 3 cycles at the lowest frequency to 10.24 at the highest and increasing linearly across this range. We extracted the event-related power spectrum changes of two-hundred points across a time range beginning 714 ms before cue onset and ending 1714 ms after the cue. As we were interested in power differences between ipsilateral and contralateral hemifields, we initially calculated raw power values with no baseline correction.

To track the deployment of attention, we calculated the difference between contralateral and ipsilateral alpha power as observed at a set of symmetrically located channels (O1/O2, PO3/PO4, PO7/PO8), subsequently computing the difference in alpha power lateralization between horizontal and vertical rectangle conditions. The specific channels used were chosen based on previous research investigating alpha oscillations in the context of attentional selection (<u>Kelly et al., 2006</u>; <u>Wang et al., 2019</u>; <u>Redding & Fiebelkorn, 2024</u>). T-values were generated for each time-frequency point against a null hypothesis of zero and results were cluster-corrected to control family-wise error. Each point with t-value greater than 1.699 (df = 29, p < 0.05, *one-sided*) became part of a cluster with the neighboring values that also met this criterion.

Cluster mass was used as the permutation statistic (<u>Maris & Oostenveld, 2007</u>). Cluster correction was applied to data observed from 300 ms (end of auditory cue) to 800 ms and from 5.9 to 40 Hz.

### ERP Analysis

ERPs were calculated with a precue baseline of 100 ms. We isolated the ADAN by calculating the difference in ipsilateral and contralateral cue-elicited ERPs as observed at two sets of four frontal channels - F3/F4, F5/F6, FC3/FC4, FC5/FC6 (Störmer, Green, & McDonald, 2009; Seiss et al., 2007). The ADAN was defined as the mean difference between these clusters in a 300 - 500 ms time window after cue onset.

## Results

### Alpha Oscillations

If attention automatically spreads along object contours, or if attention has the effect of enhancing the entirety of an attended object, we reasoned that cue-elicited alpha laterality should emerge more strongly in the vertical rectangle condition than in the horizontal rectangle condition. The results supported this hypothesis. As illustrated in Figure 3, cluster-corrected contralateral-minus-ipislateral alpha power at clusters of posterior channels was greater in the vertical rectangle condition than in the horizontal rectangle condition (one-tailed H0: true mean higher than 0, p_cluster_ = 0.038). This effect emerged around 500-600 ms after the onset of the audio spatial cue.

**Figure 3.**
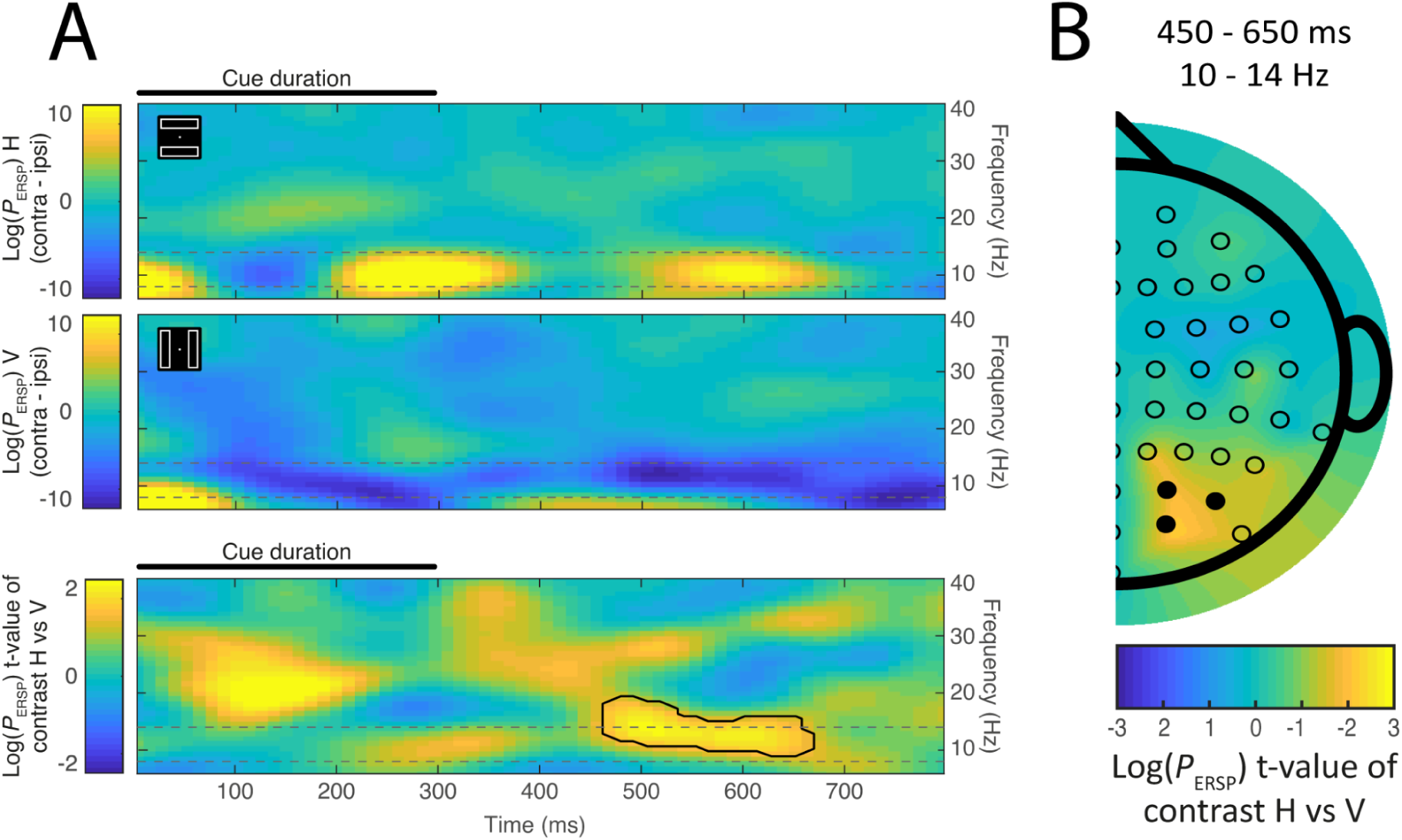
The effect of rectangle orientation on lateralized alpha amplitude. The first two rows of **Panel A** show the contralateral-minus-ipsilateral difference in time-frequency space during horizontal and vertical rectangle conditions respectively. Cool colors (blue) reflect reduced power contralateral to the cued location, while warm colors reflect greater power contralateral to the cued location. The third row illustrates the statistical contrast of data illustrated in the first two panels (horizontal-minus-vertical). Warm colors in this third row identify a combination of the increase in alpha laterality when the bars are vertical, with lower alpha power contralateral rather than ipsilateral to the cue location, and the decrease or reversal of this effect when the bars are horizontal. Results from cluster-level statistical analysis are identified; the emergence of a significant cluster (solid line) at approximately 500 - 600 ms captures the key effect. The results support the notion that the task-irrelevant rectangle is attentionally selected, and thus that attention is more strongly lateralized when the rectangle is vertical rather than horizontal. Alpha was measured as mean signal at a set of lateral occipital electrodes (O2 / PO4 / PO8, O1 / PO3 / PO7). Time zero represents the onset of the auditory cue, which lasted 300 ms. The alpha range (8-12 Hz) is identified by broken lines. For this illustration, but not statistical analyses, the time-frequency matrix was smoothed with a 2D Gaussian filter (σ = 1; filter size = 10×10). **Panel B** illustrates the scalp topography of the lateralized alpha effect depicted in the bottom row of Panel A. This half-head scalp map was created from the contralateral–ipsilateral difference waves by mirroring the data across the midline and artificially setting the values on the midline to zero (Hickey, Di Lollo, & McDonald, 2009). The topographic map reflects the emergence of the difference in contralateral-minus-ipsilateral between horizontal and vertical conditions as calculated at 8 - 12 Hz for symmetric pairs of electrodes across the scalp. Topography reflects mean signal from 450 - 650 ms after cue onset. The locations of the set of electrodes considered for the analysis is identified in the topographical plot with filled black markers. The MATLAB function *plot_topography* was used (Martínez-Cagigal, 2024).

Though we approached the experiment with an expectation of results in the alpha band, the cluster identified spanned alpha (8-14 Hz) and low beta bands (14 - 20 Hz; <u>Sassenhagen & Draschkow, 2019</u>). To test the specific involvement of alpha in object-based attention, we conducted an additional analysis, in which we band-pass filtered the experimental results to isolate the 8-14 Hz frequency band and repeated the analysis described above. This identified a significant cluster in the alpha frequency range that had much the same temporal characteristics of the alpha/beta cluster described above.

As can be noted in the last row of Figure 3A, there is a prominent conditional difference in lateralization that begins soon after onset of the auditory cue in the beta frequency band (∼12 - 30 Hz). The analyses described above were limited to the interval following offset of the auditory cue, but in separate analysis of the 0 - 300 ms interval where the auditory cue occurred, we found a significant cluster in the beta band. This effect emerges before the semantic content of the cue could be available, and its latency suggests it may reflect auditory sensory activity (Gourévitch, Martin, Postal, & Eggermont, 2020), and we do not consider it further.

### ADAN

The cue-elicited ERPs at anterior electrode locations for horizontal rectangle and vertical rectangle conditions are illustrated in Figure 4. The difference between ipsilateral and contralateral channels in the 300 - 500 ms latency range is significant from zero for vertical rectangle trials (*μ* = 0.308, bootstrapped 95%, CI = [0.054 0.635], *d* = 0.374), but not for horizontal rectangle trials (*μ* = −0.075, bootstrapped 95%, CI = [-0.293 0.160], *d* = 0.117). Notably, the contrast between contralateral and ipsilateral channels exhibits a pronounced increase for vertical rather than horizontal rectangles (*μ* = 0.383, bootstrapped 95% CI = [0.004 0.854], *d* = 0.315). This difference is highlighted in the contralateral-minus-ipsilateral difference waves illustrated in Figure 4C.

**Figure 4.**
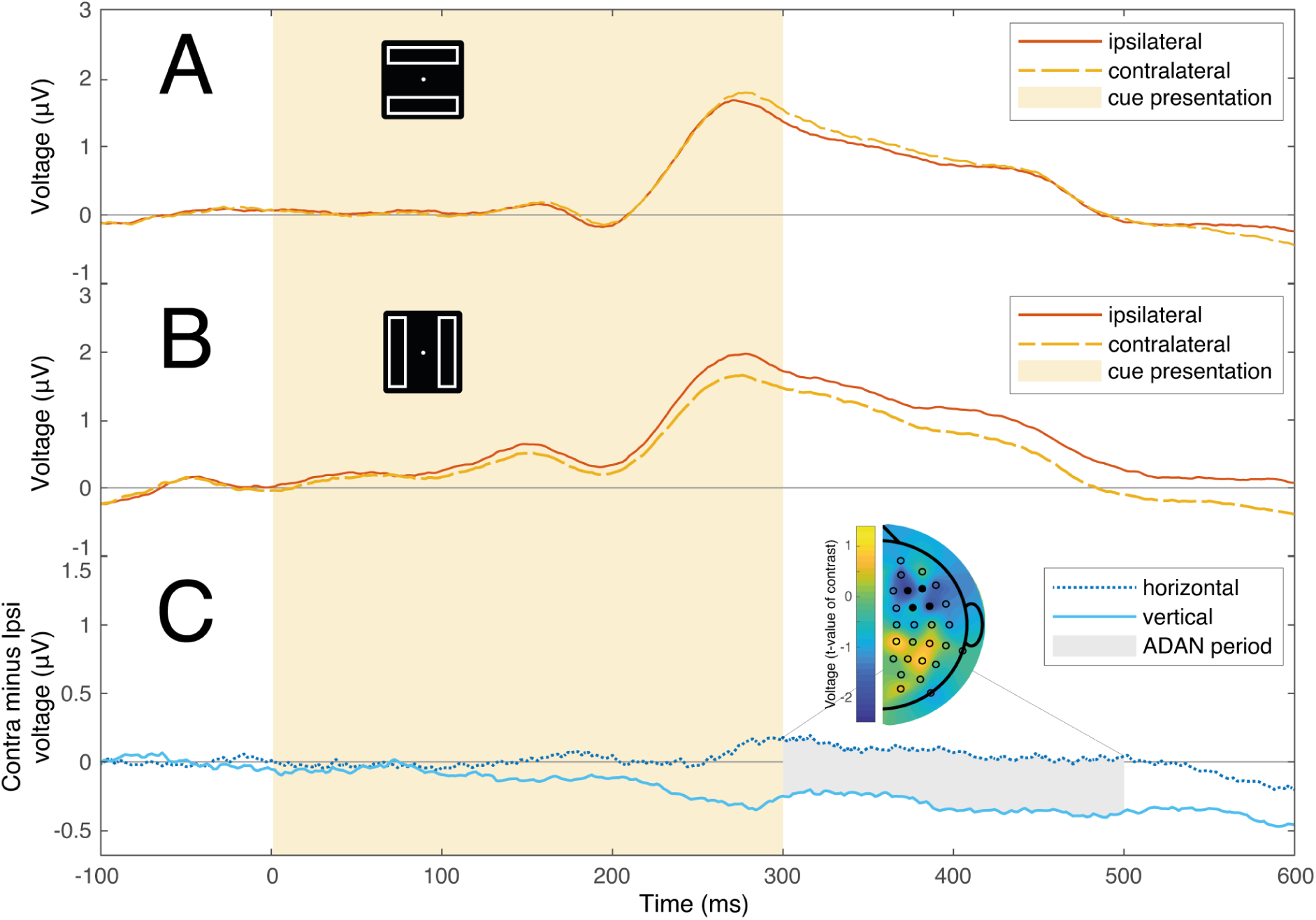
Anterior ERPs elicited by the cue stimulus. Time 0 represents the start of the auditory cue, which lasted for 300 ms (light yellow background). **Panels A and B** show ERPs measured at channels contralateral (yellow dashed line) and ipsilateral (red solid line) to the cue when the rectangles were horizontal or vertical. **Panel C** shows the difference waves between ipsilateral and contralateral ERPs in horizontal rectangle (dark blue dotted line) and vertical rectangle conditions (light blue solid line). Gray shading identified the typical interval of the ADAN (300-500ms). The topographical map (frontal scalp is at the top) shows the scalp distribution of t-values resulting from a test of the difference in lateralized signals illustrated in panels A and B (df = 29). As in Experiment 1, this topographic map reflects the contralateral-minus-ipsilateral difference as calculated for symmetric pairs of electrodes across the scalp, with values for electrodes on the vertical midline set to zero (Hickey, Di Lollo, & McDonald, 2009). The ERPs reflect mean signal at electrode clusters in the left (F3, F5, FC3, FC5) and right hemispheres (F4, F6, FC4, FC6), and the location of these electrodes is identified in the topographical plot with filled black markers.

### Behavior

Target responses were reliably faster when the target location was validly cued, as compared to conditions where the target appeared at any uncued location (591 vs. 731 ms, t(29) = 8.99, p < .001, *d* = 1.344). However, we did not find a significant difference in reaction time for invalidly cued targets when on the same-object versus a different-object (730 vs. 731 ms, t(29) = 0.36, p = .721, d = 0.013). Accuracy closely matched the reaction time results with reliably higher accuracies following valid trials (96.8% vs 86.6%, t(29) = 5.38, p < .001, d = 1.249) and no significant difference between same- and different-object conditions (86.5% vs. 86.7%, t(29) = 0.25, p = .807, d = 0.018).

The absence of any behavioural effect of object prioritization is in contrast to earlier results from Goldsmith and Yeari (2003), where similar endogenous auditory cues were employed. Note, however, that characteristics of our design differed substantially from this earlier work. In line with Abrams and Law (2000), we employed longer cue-target intervals to give participants sufficient opportunity to interpret the endogenous cues and to deploy attention. Moreover, the cue-to-target delay varied across trials to allow for the deconvolution of cue-elicited and target-elicited brain activity. The long and uncertain cue-target interval (1200-1700 ms) may have reduced the participants’ ability to maintain prioritization of the entire cued object (Lou, Lorist, & Pilz, 2023).

As described in the Introduction, recent results have shown that behavioural effects associated with object-based attention in the two-rectangle paradigm may be strongest for horizontally-oriented rectangles, sometimes only emerging in this condition (Al-Janabi & Greenberg, 2016; Chen & Cave, 2019). To test this in the current data, we separated invalidly cued trials as a function of rectangle orientation. As illustrated in Figure 5, rectangle orientation had a dramatic effect on the behavioural results. A 2-way ANOVA with factors for rectangle orientation (horizontal vs. vertical) and cue-target relationship (same-object vs. different-object) identified an interaction between rectangle orientation and cue-target relationship (F(1,29) = 35.350, p < .001, *ƞ_p_^2^* = .549). Further analysis showed that while there was a speeding of invalid target reaction time in same-object trials in the horizontal-rectangle condition (horizontal-same vs. horizontal-different: 709 vs 759 ms, t(29) = 5.04, Bonferroni-corrected p < .001, *d* = 0.340), the reverse pattern emerged in the vertical-rectangle condition (vertical-same vs. vertical-different: 757 vs 709 ms, t(29) = 5.71, Bonferroni-corrected p < .001, *d* = 0.305). Separate analysis of validly cued trials identified no effect of rectangle orientation (632 vs. 630 ms, t(29) = 0.745, p = .462, *d* = 0.020).

**Figure 5.**
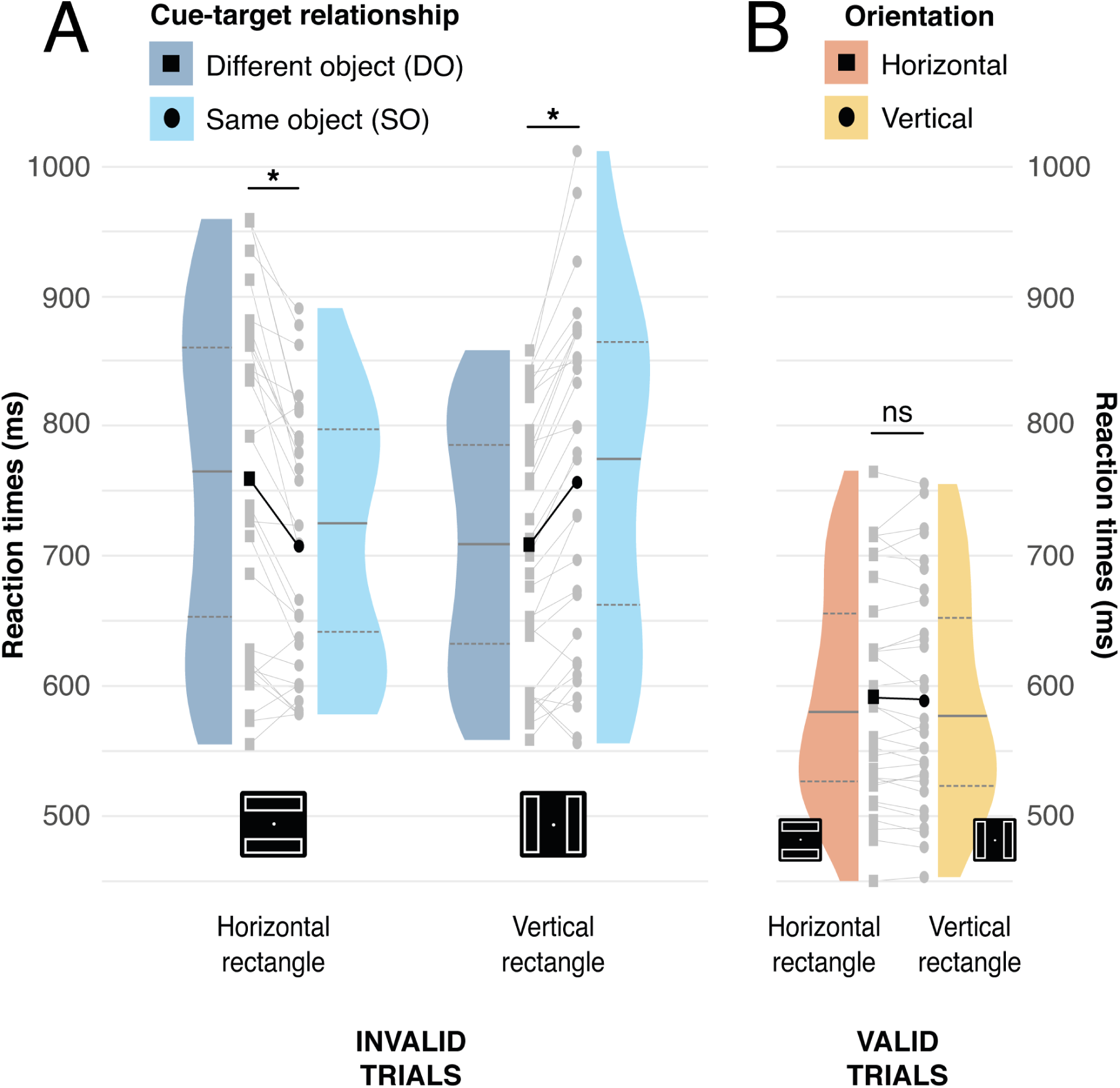
Reaction times (RT) from the behavioral task. Panel A shows Invalid Same and Different-object conditions as a function of rectangle orientation. Black squares and circles represent mean conditional reaction times for Invalid Same- and Different-object conditions, respectively. Darker and lighter blue represent invalid same- and different-object conditions, respectively. Panel B shows the valid cue condition. Black squares and circles represent mean conditional reaction times for Horizontal and Vertical conditions, respectively. Darker and lighter orange represent invalid Horizontal and Vertical conditions, respectively. In both panels, the distribution of participant mean performance is indicated for each condition, and per-participant results are illustrated in grey. The median, first, and third quartile are indicated for each distribution, and * indicate statistical significance (⍺ = .05).

Accuracy showed a similar pattern. A 2-way ANOVA with factors for rectangle orientation and cue-target relationship identified only a significant interaction effect (F(1,29) = 31.394, p < .001, *ƞ_p_^2^* = 0.520). Pairwise comparison highlighted a significant difference in opposite directions: in the horizontal condition, accuracy was greater for same-objects trials (horizontal-same vs. horizontal-different: 92.0% vs 81.5%, t(29) = 5.029, Bonferroni-corrected p < .001, *d* = 0.798), but this effect reversed in the vertical condition (vertical-same vs. vertical-different: 80.6% vs 91.5%, t(29) = −5.303, Bonferroni-corrected p < .001, *d* = 0.892).

## Discussion

A substantial body of research suggests that selective attention is influenced by the presence of visual objects. Specifically, when attention is deployed to one location on an object, other locations on the same item appear to be prioritized (Egly, Driver, & Rafal, 1994; Moore, Yantis, & Vaughan, 1998; Abrams & Law, 2000; Chen & Cave, 2008). However, recent findings have raised challenges to this view. First, behavioural evidence of object prioritization is stronger when objects span the visual hemifields, and in some cases only emerges in this scenario (Al-Janabi & Greenberg, 2016; Chen & Cave, 2019; Francis & Thunnell, 2022; Pilz, Roggeveen, Creighton, Bennett, & Sekuler, 2012). This has led to speculation that putative object-based attention may in fact result from the presence of independent attention resources in each cerebral hemisphere (Luck et al., 1989), which facilitates shifts of attention across the vertical meridian (Barnas & Greenberg, 2016b, 2024). As a result, evidence of object prioritization may be confounded with effects linked to the independence of attentional systems in the two visual cortices. Second, manipulations of object status have concomitant and often unacknowledged effects on low-level stimuli characteristics, such as creating visual clutter between cue and target locations (Rosenholtz, 2024). This may create a small cost to perceptual resolution of the target similar to that observed in studies of visual crowding (Whitney & Levi, 2011; Chen, Cave, Basu, Suresh, & Wiltshire, 2020).

To address these issues, we used EEG to directly index the deployment of attention. EEG has been used to study object-based attention before, but existing studies have concentrated on how objects impact the sensory processing of targets. This work has shown that invalidly-cued targets that appear on cued objects elicit a larger posterior N1 ERP component (Martínez et al., 2006) that is similar (but not identical; He, Fan, Zhou, & Chen, 2004; He, Humphreys, Fan, Chen, & Han, 2008) to that observed when targets are validly cued (Mangun & Hillyard, 1991;Handy & Mangun, 2000). In contrast to this work, we focus on EEG and ERP effects in the interval immediately following the cue and prior to appearance of the target (e.g., Harter, Miller, Price, LaLonde & Keyes, 1989; Worden, Foxe, Wang & Simpson, 2000).

There are two prominent findings in our results. First, participants used the endogenous cue to guide the deployment of spatial attention. This is evident in the cue-elicited emergence of posterior EEG alpha lateralization, the presence of the evoked anterior ADAN ERP component, and the significant effect of cue validity on behavioural responses to the targets. Second, and critically, we find that the hemispheric pattern of alpha and ADAN is impacted by the orientation of the task-irrelevant rectangles. These lateralized EEG/ERP effects are more pronounced when a cued rectangle is oriented vertically, and thus appears entirely within one visual hemifield, than when a cued rectangle is oriented horizontally, and therefore spans the vertical meridian of the visual field. This supports the idea that attentional prioritization of objects leads to greater lateralization of attention when the attended object is in one visual hemifield (and thus one hemisphere), compared to when the object spans the vertical meridian and involves both hemispheres.

Our findings provide direct evidence that object-based attention is a genuine psychological and neural phenomenon. However, this does not mean that concerns regarding the confounding impact of hemifield anisotropies or visual crowding on responses to the target are misplaced. We do not find the hallmark same-object benefit on manual response to the target. Instead, we find a strong anisotropy effect: participants are faster to respond to targets on the cued object when the rectangles are oriented horizontally, with the reverse emerging when rectangles are oriented vertically. This mirrors earlier findings from Pilz et al. (2012, Exp. 2) and suggests that attentional deployment across the hemifields facilitates lateralization of attention, regardless of the presence or structure of visual objects.

There is thus disparity between our behavioural results, which show no evidence of object prioritization, and our EEG/ERP results, which do. One account for this is that the object prioritization we see in EEG/ERP is not sustained through the long and variable cue-target interval we employed (Lou, Lorist, & Pilz, 2023). However, this interpretation has an important caveat. Pilz et al. (2012, Exp 2) also observed a strong hemifield anisotropy effect in the two-rectangle paradigm, without any evidence of object prioritization, though the cue-target intervals employed in that study were short and consistent. This motivates an alternative account, namely that object prioritization emerges and persists until appearance of the target, but that this effect is somehow negated or overshadowed when a more robust effect of hemifield anisotropy emerges. There is the clear opportunity for further research to identify why strong effects of hemifield anisotropy emerge in some experiments employing the two-rectangle task (eg. Pilz et al., 2012; Barnas & Greenberg, 2024), but not in others (eg. Egly, Driver, & Rafal, 1994), and if the emergence of hemifield anisotropy is a predictor of the absence of object prioritization at the time of target onset.

If it proves to be the case that long cue-target duration reduces the effect of object prioritization on behaviour, this suggests that participants may strategically disengage from the cued rectangle when they are given sufficient time to do so, and this raises a broader issue regarding the role of strategy in object-based attention. The notion that object prioritization is necessary and automatic has been challenged by results suggesting a strategic basis for the effect (Shomstein, 2012; Shomstein & Yantis, 2002; Shomstein & Johnson, 2013). For example, object prioritization in the two-rectangle paradigm disappears when the target location is known in advance (Drummond & Shomstein, 2010a) or when other strategies have greater economic utility (Shomstein & Johnson, 2013; though see Grignolio, Acunzo, & Hickey, 2024; Diao et al., 2024). Our results from the ADAN ERP component are relevant in this context. In contrast to lateral alpha, which is thought to reflect low-level, mechanistic effects of inhibitory gating in sensory cortex (Jensen & Mazaheri, 2010; Jensen, 2024), the ADAN is thought to reflect anterior brain structures - possibly including the lateral frontal eye fields - that are involved in the strategic control of attention (Eimer, van Velzen, & Forster, 2003; Hopf & Mangun, 2000). We find that object prioritization is expressed in ADAN lateralization, and this suggests that such prioritization is represented in high-level, anterior cortical structures responsible for the strategic control of selection. While far from conclusive, this is in line with accumulating evidence that object-based attention reflects a strategic approach to task completion (Shomstein, 2012).

Our results are consistent with two models of object-based attention proposed in the literature. One model, often implicitly adopted in the text above, suggests that attention is deployed to a spatial location and then spreads along object contours (possibly to support definition of the visual object; Duncan, 1994). The alternative is that objects may be defined in the visual system prior to the deployment of attention, with attentional deployment to a location intrinsically linked with attentional selection of objects at that location (Scholl, 2001). Though the current results do not allow us to differentiate between these possibilities, recent results from fMRI provide compelling evidence of attentional spreading. Ekman, Roelfsema, and de Lange (2020) used advanced techniques to map the effect of a spatial cue on the representation of an underlying object in V1, showing a spread of activity from populations of neurons representing the initially selected region of the object to others representing the rest of the object.

In conclusion, our study demonstrates that selective attention is influenced by the presence of irrelevant visual objects. We directly index this effect using EEG, finding that lateralized alpha and the ADAN component of the ERP vary as a function of the orientation of irrelevant rectangle stimuli. Critically, we index the influence of these irrelevant objects on attention in the interval preceding appearance of the target, and our results are therefore unaffected by some potential confounds that have clouded interpretation of behavioural studies of object-based attention.

## Acknowledgements

Our thanks to Carmel Mevorach for support with student mobility and the Alan Turing Institute for travel funding to DG, and to members of the Center for Mind and Brain at UC Davis for their support of this work. This study was supported in part by NIMH grant MH117991 to GRM. DG and CH are supported by the European Research Council (ERC) under the European Union Horizon 2020 Research and Innovation Program (Grant Agreement 804360 to CH).

## References

Abrams, R. A., & Law, M. B. (2000). Object-based visual attention with endogenous orienting. Perception & Psychophysics, 62(4), 818–833.

Acunzo, D. J., MacKenzie, G., & van Rossum, M. C. (2012). Systematic biases in early ERP and ERF components as a result of high-pass filtering. Journal of neuroscience methods, 209(1), 212–218.

Al-Janabi, S., & Greenberg, A. S. (2016). Target–object integration, attention distribution, and object orientation interactively modulate object-based selection. Attention, Perception, & Psychophysics, 78, 1968–1984.

Audacity Team. (2021). Audacity (Version 3.2.4) [Computer software]. https://www.audacityteam.org/

Barnas, A. J., & Greenberg, A. S. (2016b). Visual field meridians modulate the reallocation of object-based attention. Attention, Perception, & Psychophysics, 78, 1985–1997.

Barnas, A. J., & Greenberg, A. S. (2024). The object-based shift direction anisotropy is modulated by the horizontal visual field meridian. Quarterly Journal of Experimental Psychology, 17470218241230988.

Bell, A. J., & Sejnowski, T. J. (1995). An information-maximization approach to blind separation and blind deconvolution. Neural computation, 7(6), 1129–1159.

Carrasco, M., Giordano, A. M., & McElree, B. (2004). Temporal performance fields: Visual and attentional factors. Vision research, 44(12), 1351–1365.

Cavanagh, P., Caplovitz, G. P., Lytchenko, T. K., Maechler, M. R., Tse, P. U., & Sheinberg, D. L. (2023). The architecture of object-based attention. Psychonomic Bulletin & Review, 30(5), 1643–1667.

Chen, Z. (2012). Object-based attention: A tutorial review. Attention, Perception, & Psychophysics, 74, 784–802.

Chen, Z., & Cave, K. R. (2008). Object-based attention with endogenous cuing and positional certainty. Perception & Psychophysics, 70, 1435–1443.

Chen, Z., & Cave, K. R. (2019). When is object-based attention not based on objects?. Journal of experimental psychology: human perception and performance, 45(8), 1062.

Chen, Z., Cave, K. R., Basu, D., Suresh, S., & Wiltshire, J. (2020). A region complexity effect masquerading as object-based attention. Journal of Vision, 20(7), 24–24.

Cohen, E. H., & Tong, F. (2015). Neural mechanisms of object-based attention. Cerebral Cortex, 25(4), 1080–1092.

Corballis, M. C., & Roldan, C. E. (1975). Detection of symmetry as a function of angular orientation. Journal of Experimental Psychology: Human Perception and Performance, 1(3), 221.

Corbett, J. E., & Carrasco, M. (2011). Visual performance fields: Frames of reference. PLoS One, 6(9), e24470.

Coslett, H. B., & Saffran, E. (1991). Simultanagnosia: To see but not two see. Brain, 114(4), 1523–1545.

Davis, G., & Driver, J. (1997). Spreading of visual attention to modally versus amodally completed regions. Psychological Science, 8(4), 275–281.

Delorme, A., & Makeig, S. (2004). EEGLAB: an open source toolbox for analysis of single-trial EEG dynamics including independent component analysis. Journal of neuroscience methods, 134(1), 9–21.

Diao, F., Hu, X., Zhang, T., Gao, Y., Zhou, J., Kong, F., & Zhao, J. (2024). The Impact of Reward Object on Object-Based Attention. Behavioral Sciences, 14(6), 505.

Drummond, L., & Shomstein, S. (2010a). Object-based attention: Shifting or uncertainty?. Attention, Perception, & Psychophysics, 72, 1743–1755.

Duncan, J. (1984). Selective attention and the organization of visual information. Journal of experimental psychology: General, 113(4), 501.

Egly, R., Driver, J., & Rafal, R. D. (1994). Shifting visual attention between objects and locations: evidence from normal and parietal lesion subjects. Journal of Experimental Psychology: General, 123(2), 161.

Eimer, M., Velzen, J. V., & Driver, J. (2002). Cross-modal interactions between audition, touch, and vision in endogenous spatial attention: ERP evidence on preparatory states and sensory modulations. Journal of cognitive neuroscience, 14(2), 254–271.

Eimer, M., Forster, B., & Van Velzen, J. (2003). Anterior and posterior attentional control systems use different spatial reference frames: ERP evidence from covert tactile-spatial orienting. Psychophysiology, 40(6), 924–933.

Ekman, M., Roelfsema, P. R., & de Lange, F. P. (2020). Object selection by automatic spreading of top-down attentional signals in V1. Journal of Neuroscience, 40(48), 9250–9259.

Foster, J. J., & Awh, E. (2019). The role of alpha oscillations in spatial attention: limited evidence for a suppression account. Current opinion in psychology, 29, 34–40.

Foxe, J. J., & Snyder, A. C. (2011). The role of alpha-band brain oscillations as a sensory suppression mechanism during selective attention. Frontiers in psychology, 2, 154.

Francis, G., & Thunell, E. (2022). Excess success in articles on object-based attention. Attention, Perception, & Psychophysics, 84(3), 700–714.

Goldsmith, M., & Yeari, M. (2003). Modulation of object-based attention by spatial focus under endogenous and exogenous orienting. Journal of Experimental Psychology: Human Perception and Performance, 29(5), 897.

Gourévitch, B., Martin, C., Postal, O., & Eggermont, J. J. (2020). Oscillations in the auditory system and their possible role. Neuroscience & Biobehavioral Reviews, 113, 507–528.

Grignolio, D., Acunzo, D. J., & Hickey, C. (2024). Object-based attention is accentuated by object reward association. Journal of Experimental Psychology: Human Perception and Performance, 50(3), 280.

Handy, T.C., & Mangun, G.R. (2000). Attention and spatial selection: Electrophysiological evidence for modulation by perceptual load. Perception and Psychophysics, 62, 175–186.

Harter, M. R., Miller, S. L., Price, N. J., LaLonde, M. E., & Keyes, A. L. (1989). Neural processes involved in directing attention. Journal of Cognitive Neuroscience, 1, 223–237.

He, X., Fan, S., Zhou, K., & Chen, L. (2004). Cue validity and object-based attention. Journal of Cognitive Neuroscience, 16(6), 1085–1097.

He, X., Humphreys, G., Fan, S., Chen, L., & Han, S. (2008). Differentiating spatial and object-based effects on attention: An event-related brain potential study with peripheral cueing. Brain Research, 1245, 116–125.

Hickey, C., Di Lollo, V., & McDonald, J. J. (2009). Electrophysiological indices of target and distractor processing in visual search. Journal of Cognitive Neuroscience, 21(4), 760–775.

Holmes, A., Mogg, K., Garcia, L. M., & Bradley, B. P. (2010). Neural activity associated with attention orienting triggered by gaze cues: A study of lateralized ERPs. Social neuroscience, 5(3), 285–295.

Hopf, J. M., & Mangun, G. R. (2000). Shifting visual attention in space: an electrophysiological analysis using high spatial resolution mapping. Clinical neurophysiology, 111(7), 1241–1257.

Jensen, O., & Mazaheri, A. (2010). Shaping functional architecture by oscillatory alpha activity: gating by inhibition. Frontiers in human neuroscience, 4, 186.

Jensen, O. (2024). Distractor inhibition by alpha oscillations is controlled by an indirect mechanism governed by goal-relevant information. Communications Psychology, 2(1), 36.

Kelly, S. P., Lalor, E. C., Reilly, R. B., & Foxe, J. J. (2006). Increases in alpha oscillatory power reflect an active retinotopic mechanism for distracter suppression during sustained visuospatial attention. Journal of neurophysiology, 95(6), 3844–3851.

Klimesch, W., Sauseng, P., & Hanslmayr, S. (2007). EEG alpha oscillations: the inhibition–timing hypothesis. Brain research reviews, 53(1), 63–88.

Lou, H., Lorist, M. M., & Pilz, K. S. (2023). Variability in the temporal dynamics of object-based attentional selection. Plos one, 18(11), e0294252.

Luria, A. R. (1959). Disorders of “ldquo; simultaneous perception” in a case of bilateral occipito-parietal brain injury. Brain, 82(3), 437–449.

Luck, S. J., Hillyard, S. A., Mangun, G. R., & Gazzaniga, M. S. (1989). Independent hemispheric attentional systems mediate visual search in split-brain patients. Nature, 342(6249), 543–545.

Mangun, G. R., & Hillyard, S. A. (1991). Modulations of sensory-evoked brain potentials indicate changes in perceptual processing during visual-spatial priming. Journal of Experimental Psychology: Human perception and performance, 17(4), 1057.

Maris, E., & Oostenveld, R. (2007). Nonparametric statistical testing of EEG-and MEG-data. Journal of neuroscience methods, 164(1), 177–190.

Moore, C. M., Yantis, S., & Vaughan, B. (1998). Object-based visual selection: Evidence from perceptual completion. Psychological science, 9(2), 104–110.

Nobre, A. C., Gitelman, D. R., Dias, E. C., & Mesulam, M. M. (2000). Covert visual spatial orienting and saccades: overlapping neural systems. Neuroimage, 11(3), 210–216.

Oostenveld, R., & Praamstra, P. (2001). The five percent electrode system for high-resolution EEG and ERP measurements. Clinical neurophysiology, 112(4), 713–719.

Pilz, K. S., Roggeveen, A. B., Creighton, S. E., Bennett, P. J., & Sekuler, A. B. (2012). How prevalent is object-based attention?. PloS one, 7(2), e30693.

Pion-Tonachini, L., Kreutz-Delgado, K., & Makeig, S. (2019). ICLabel: An automated electroencephalographic independent component classifier, dataset, and website. NeuroImage, 198, 181–197.

Popov, T., Gips, B., Kastner, S., & Jensen, O. (2019). Spatial specificity of alpha oscillations in the human visual system. Human Brain Mapping, 40(15), 4432–4440.

Praamstra, P., Boutsen, L., & Humphreys, G. W. (2005). Frontoparietal control of spatial attention and motor intention in human EEG. Journal of neurophysiology, 94(1), 764–774.

Redding, Z. V., & Fiebelkorn, I. C. (2024). Separate cue-and alpha-related mechanisms for distractor suppression. Journal of neuroscience, 44(25).

Reppa, I., Schmidt, W. C., & Leek, E. C. (2012). Successes and failures in producing attentional object-based cueing effects. Attention, Perception, & Psychophysics, 74, 43–69.

Rosenholtz, R. (2024). Visual Attention in Crisis. Behavioral and Brain Sciences, 1-32.

Sassenhagen, J., & Draschkow, D. (2019). Cluster-based permutation tests of MEG/EEG data do not establish significance of effect latency or location. Psychophysiology, 56(6), e13335.

Scholl, B. J. (2001). Objects and attention: The state of the art. Cognition, 80(1-2), 1–46.

Seiss, E., Gherri, E., Eardley, A. F., & Eimer, M. (2007). Do ERP components triggered during attentional orienting represent supramodal attentional control?. Psychophysiology, 44(6), 987–990.

Shomstein, S. (2012). Object-based attention: strategy versus automaticity. Wiley Interdisciplinary Reviews: Cognitive Science, 3(2), 163–169.

Shomstein, S., & Johnson, J. (2013). Shaping attention with reward: Effects of reward on space-and object-based selection. Psychological science, 24(12), 2369–2378.

Shomstein, S., & Yantis, S. (2002). Object-based attention: Sensory modulation or priority setting?. Perception & psychophysics, 64(1), 41–51.

Stoermer, V. S., Green, J. J., & McDonald, J. J. (2009). Tracking the voluntary control of auditory spatial attention with event-related brain potentials. Psychophysiology, 46(2), 357–366.

Van Velzen, J., Forster, B., & Eimer, M. (2002). Temporal dynamics of lateralized ERP components elicited during endogenous attentional shifts to relevant tactile events. Psychophysiology, 39(6), 874–878.

Martínez-Cagigal, V. (2024). Topographic EEG/MEG plot (https://www.mathworks.com/matlabcentral/fileexchange/72729-topographic-eeg-meg-plot), MATLAB Central File Exchange. Retrieved November 29, 2024.

Walker, R. (1995). Spatial and object-based neglect. Neurocase, 1(4), 371–383.

Wang, B., van Driel, J., Ort, E., & Theeuwes, J. (2019). Anticipatory distractor suppression elicited by statistical regularities in visual search. Journal of cognitive neuroscience, 31(10), 1535–1548.

Watson, S. E., & Kramer, A. F. (1999). Object-based visual selective attention and perceptual organization. Perception & Psychophysics, 61(1), 31–49.

Whitney, D., & Levi, D. M. (2011). Visual crowding: A fundamental limit on conscious perception and object recognition. Trends in cognitive sciences, 15(4), 160–168.

Worden, M. S., Foxe, J. J., Wang, N., & Simpson, G. V. (2000). Anticipatory biasing of visuospatial attention indexed by retinotopically specific alpha-band electroencephalography increases over occipital cortex. The Journal of neuroscience: the official journal of the Society for Neuroscience, 20(6), RC63–RC63.

Yamaguchi, S., Tsuchiya, H., & Kobayashi, S. (1995b). Electrophysiologic correlates of visuo-spatial attention shift. Electroencephalography and clinical Neurophysiology, 94(6), 450–461.

Zani, A., Crotti, N., Marzorati, M., Senerchia, A., & Proverbio, A. M. (2023). Acute hypoxia alters visuospatial attention orienting: an electrical neuroimaging study. Scientific Reports, 13(1), 22746.

